# A Modified ACE2 peptide mimic to block SARS-CoV2 entry

**DOI:** 10.1101/2020.05.07.082230

**Authors:** Suman Saurabh, Shubh Sanket Purohit

## Abstract

A 23-residue peptide fragment that forms a part of the *α*-1 helix of the ACE2 peptidase domain, the recognition domain for SARS-CoV2 on the ACE2 receptor, holds the potential as a drug to block the viral receptor binding domain (RBD) from forming a complex with ACE2. The peptide has recently been shown to bind the viral RBD with good efficiency. Here, we present a detailed analysis of the energetics of binding of the peptide to the SARS-CoV2 RBD. We use equilibrium molecular dynamics simulation to study the dynamics of the complex. We perform end-state binding energy calculations to gain a residue-level insight into the binding process and use the information to incorporate point mutations into the peptide. We demonstrate using binding energy calculations that the peptide with certain point mutations, especially E17L, shows a stronger binding to the RBD as compared to the wild type peptide. We propose that the modified peptide will thus be more efficient in blocking RBD-ACE2 binding.

## Introduction

Since December 2019 after the initial cases of Covid-19 were detected and SARS-CoV2, a virus closely related to the severe acute respiratory syndrome coronavirus (SARS-CoV), was recognised as the virus underlying the disease,^1^ laboratories around the world have been working towards finding a counter. With the declaration of Covid-19 as a public health emergency, serious efforts to develop a cure gained speed. Various strategies have been put to work for this purpose. As the development of a new vaccine from the scratch requires a very long time, the rapidly growing affect of the pandemic has made researchers look into already available drugs that would be effective against SARS-CoV2 and can be alterered to increase their effectiveness against the virus.

To achieve this, one needs to have a firm understanding of the structural details of the viral RBD, the binding interface of the complex between the RBD and the human cell proteins that are involved in the process of the viral entry into the cell and the viral proteins that are involved in the successive replication process. Researchers have made impressively quick inroads towards understanding the structure of the proteins involved. Wrapp *et al.*^2^ developed a Cryo EM structure of the viral spike trimer, both in the receptor-binding (open) and closed conformations at 3.5 *Å* resolution (pdb id: 6VSB). It has been shown that SARS-CoV2 uses the same receptor, human angeotensin converting enzyme (ACE2), as that used by SARS-CoV to gain entry into the human cell.^3–5^ To make an informed choice of a molecule among the available drugs that can block the RBD-ACE2 binding requires a visual understanding of the structure and composition of the binding interface of the complex between the two proteins. Many crystal structures of the bound state of the SARS-CoV2 RBD with the ACE2 receptor have been determined. For example, Yan *et al.*^6^ determined the Cryo-EM structure of RBD-ACE2 complex in the presence of B^0^AT1 at a resolution of 3.5 *Å* at the binding interface and an overall resolution of 2.9 *Å* (pdb id: 6M17). Shang *et al.*^7^ determined the crystal structure of the ACE2-RBD complex (pdb id: 6VW1). They found that the ACE2-binding ridge in SARS-CoV-2 RBD is more compact as compared to that of SARS-CoV and multiple favourable residue changes vis-à-vis SARS-CoV lead to a stronger binding of the SARS-CoV2 RBD to ACE2.

On observing the crystal structures discussed above it is clear that unlike many other virions such as the HIV, where the receptor binding region of the spike protein is a compact hemispherical domain,^8,9^ the SARS-CoV2 RBD has an extended linear ridge-like binding region. A small-molecule SARS-CoV2 entry-inhibitor that would target the RBD would be incapable of engaging the complete ridge and would also be prone to competitive replacement. In situations like these, an extended molecule, that would be able to engage most of the residues belonging to the receptor binding region and is also rigid enough to present itself as an entropically favourable binding partner, is more suitable. Peptides present such an alternative. Most peptide drugs in use act by presenting themselves as mimics of receptor proteins involved in biological pathways.^10^ The crystal structures show that the SARS-CoV2 RBD binds ACE2 by attaching to a *α*-helical region (*α*-1 helix) in its peptidase domain (see fig.1). The RBD holds on to ACE2 such that its receptor binding ridge engages this helix, with some surrounding loops on the two proteins providing an extra reinforcement to the binding. This helical peptide fragment can thus be extracted and used, potentially, as an ACE2 mimic to block the RBD-ACE2 binding. The peptide would not just be engaging the whole binding ridge on the RBD but would also be naturally biocompatible. In a recent study by Zhang *et al.*^11^ this peptide has been proposed as a possible drug against SARS-CoV2 and its capability to bind the viral RBD has also been demonstrated. The energies involved in the binding of the isolated peptide to the viral RBD is expected to be close to that of the the RBD-ACE2 complex thus raising the possibility of the peptide being competitively replaced by ACE2 in a practical scenario. Thus, ways to modify it such that it binds stronger to the RBD need to be considered. In this work we dwell into this aspect and propose ways to strengthen the peptide-ACE2 binding.

**Figure 1:**
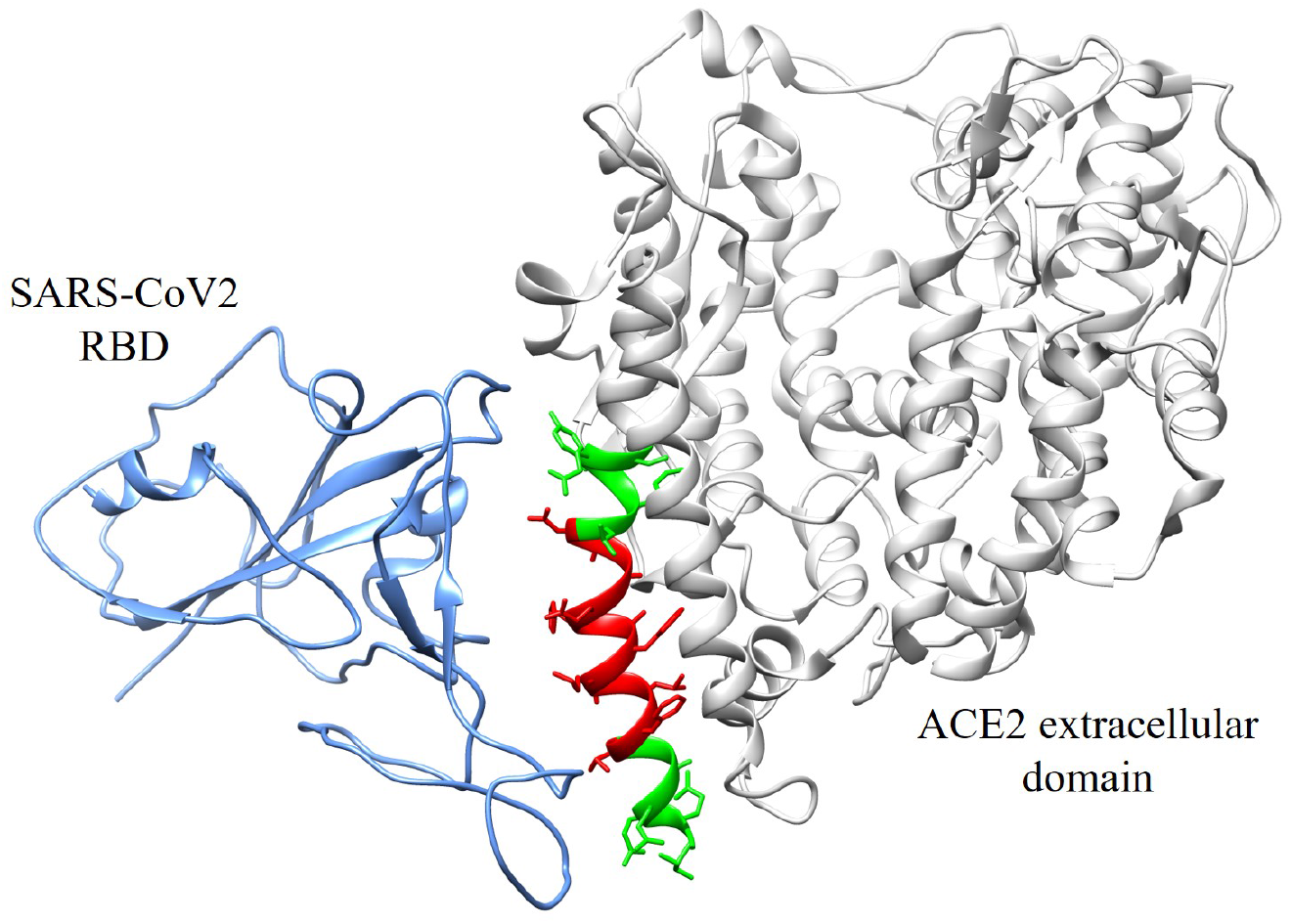
The complex between SARS-CoV2 RBD (blue) and the extra-cellular domain of ACE2 (gray), taken from the crystal structure with pdb id: 6M17.^6^ The peptide that forms a part of the viral binding region of ACE2 is shown in green. We simulate,in this work, a complex between RBD and this peptide. A subdomain of this peptide shown in red, that incorporates many key interactions is also simulated.

In recent times, molecular dynamics (MD) simulation has played an important role in the field of drug discovery. In addition to providing an atomic level resolution, it helps one observe transition pathways and important molecular motions. Although drug discovery is predominantly an experimental process, computer simulations have played an important role in guiding the decision making involved in the process and sped up the process of developing new drug molecules. With the recent increase in computational power it has been possible to perform even *μ*s to ms long MD simulations, making it possible to accurately predict the binding site of drug molecules to their targets. For example, Shan *et al.*,^12^ from multiple 100 *μ*s long simulation of systems consisting of Src kinase with the cancer drug dasatinib randomly placed inside the simulation box, succeded in recognizing the crystallographic binding site of the drugs and could also resolve the location of waters in the X-ray structure of Src-PPI complex.

Here, we perform a set of MD simulations aimed at gaining an in-depth understanding of the interaction between the SARS-CoV2 RBD and the ACE2-derived *α*-helical peptide. With the motive of modifying the peptide so as to increase its affinity towards the RBD, we study the interaction of some variants of this peptide with the RBD and demonstrate that certain point mutations can increase the binding energy of the peptide to the RBD and it can thus block RBD-ACE2 binding with higher efficiency.

## Methods

### Building the systems for simulation

The starting structure of the peptide-RBD complex was obtained by removing regions other than a single copy of the peptide and the RBD from the crystal structure with pdb id: 6M17. ^6^ The structure was enclosed in a water box of size 85*Å* × 85*Å* × 92*Å* using the xleap module of AmberTools18.^13^ The system was neutralized by adding 2 Na^+^ ions. A salt concentration of 150 mM was achieved by adding 48 Na^+^ ions and an equal number of Cl^−^ ions. TIP3P model was used for the water. Forcefield ff99sb-*ildn* ^14,15^ was used to represent the inter- and intra-molecular interactions of the protein atoms. The ions were described using the Joung Cheatham parameter set. ^16^ In the rest of the article this system with the wild-type peptide is referred to as the WT system (see table 1).

**Table 1:** Systems simulated in this study. The last 3 systems feature mutated peptides. The points of mutation have been mentioned in red in the peptide sequence.

| S.no | Simulation Model | Peptide Sequence |
| --- | --- | --- |
| 1 | Wild type (WT) | **IEEQAKTFLDKFNHEAEDLFYQS** |
| 2 | WT-S | **TFLDKFNHEAED** |
| 3 | E17L | **IEEQAKTFLDKFNHEAredLDLFYQS** |
| 4 | D18L | **IEEQAKTFLDKFNHEAredLLFYQS** |
| 5 | N13E-E17L | **IEEQAKTFLDKFredEHEAredLDLFYQS** |

In addition to the above simulation we also performed a simulation of the RBD in complex with a smaller peptide that was obtained by removing the first 6 and last 5 residues of the original 23 amino acid peptide leaving only the 12-residue central region of the peptide. In a recent study this small peptide was found to not bind to the RBD inspite of engaging in key contacts with the RBD.^11^ The system was solvated and neutralized as described above. 45 Na^+^ and an equal number of Cl^−^ ions were needed to achieve a salt concentration of 150 mM. In the rest of the article this system is referred to as WT-S (see table 1).

In order to understand the effect of certain amino acid replacements on the binding energy of the peptide-RBD complex, three different systems, referred to as E17L, D18L and N13E-E17L in the rest of the article, were built by replacing certain residues of the 23-residue peptide with their mutation partners using *Swiss PDB Viewer*.^17^ The mutation partners were decided upon based on atomistic analysis of the 6M17^6^ binding interface (described in the Results and Discussion section). Rotamers of the mutated amino acids were chosen such that there are no bad atomic contacts in the mutated peptide. After mutation the peptides were aligned with the wild type peptide in 6M17^6^ to build the RBD-peptide complex. Following this the systems were neutralised and solvated in a 150 mM NaCl environment as described above for the wild type system.

A list of all the systems and the corresponding peptide sequences is given in table 1.

### MD simulation protocol

The systems were minimized using 3000 steps of steepest descent method with the solute atoms held at their initial positions by a harmonic potential with a force constant of 500 *kcal/mol/Å*^2^. This was followed by 5000 steps of conjugate gradient. During these 5000 steps the harmonic restraint was reduced from 50 kcal/mol/*Å*^2^ to 0 in steps of 10 per 1000 steps. After minimization, the systems were heated from 0 K to 300 K within 40 ps. The systems at 300K was subjected to 1 ns of equilibration in an NPT ensemble. This was followed by a 300 ns long production run in the NVT ensemble. SHAKE constraints^18^ were used for all bonds containing hydrogen allowing the use of a time step of 2 fs. Langevin thermostat^19^ with a collision frequency of 2 ps^−1^ was used for temperature regulation while PME was used for the calculation of long range electrostatic interaction. All simulations were performed using the AMBER18 simulation package. ^13^

### End-state binding energy calculation

We used the MMGBSA method employed in the MMPBSA.py^20^ module of AMBER18^13^ to calculate the binding free energy of various RBD-peptide complexes. The method computes the binding energy (E) as: E_*bind*_ = E_*ele*_ + E_*vdw*_ + E_*int*_ + E_*sol*_ consists of changes in the electrostatic energy, E_*ele*_,van der Waals energy, E_*vdw*_, the internal energy from bonded terms, E_*int*_, and the solvent contribution, E_*sol*_. All but the last term are calculated from the force field parameters from the gas phase. The last contribution, E_*sol*_ = E_*es*_ +E_*nes*_, is the sum of the electrostatic energy of solvation, E_*es*_, calculated using the Generalized Born (GB) method, and the non-electrostatic energy of solvation, E_*nes*_, given by *γ*SASA+*β*, (*γ* = 0.00542 kcal/*Å*^2^ is the surface tension, *β* = 0.92 kcal/mol, and SASA is the solvent-accessible surface area of the molecule).

### Calculation of peptide-RBD contacts

A contact was defined to exist between a peptide residue and the RBD if any atom of the RBD fell within 3*Å* of any atom belonging to the peptide residue.

## Results and discussion

We observe, from the WT simulation, that the RBD-peptide complex stays stable throughout the 300 ns long simulation. The RMSD for the complex flattens out after around 50 ns and stays flat throughout (see fig.2). In contrast to WT, for the WT-S system, we find that the smaller 12-residue non-terminal section of the peptide (see fig.1) does not stay attached to the RBD. The time series of the distance between the centers of mass of the RBD and the peptide for the WT and the WT-S systems is shown in figure 3. One observes that while the distance for the WT system stays around 24 *Å* throughout the simulation, the small peptide detaches from the RBD and flys apart. Zhang *et al.*^11^ saw a similar behaviour in their simulations. Their experimental data also show that the smaller peptide does not bind to the RBD. We also observe that the peptide in the WT-S system also loses its helical structure (see fig.3). A loss of helical structure and reduction in rigidity would render the bound state entropically unfavourable.

**Figure 2:**
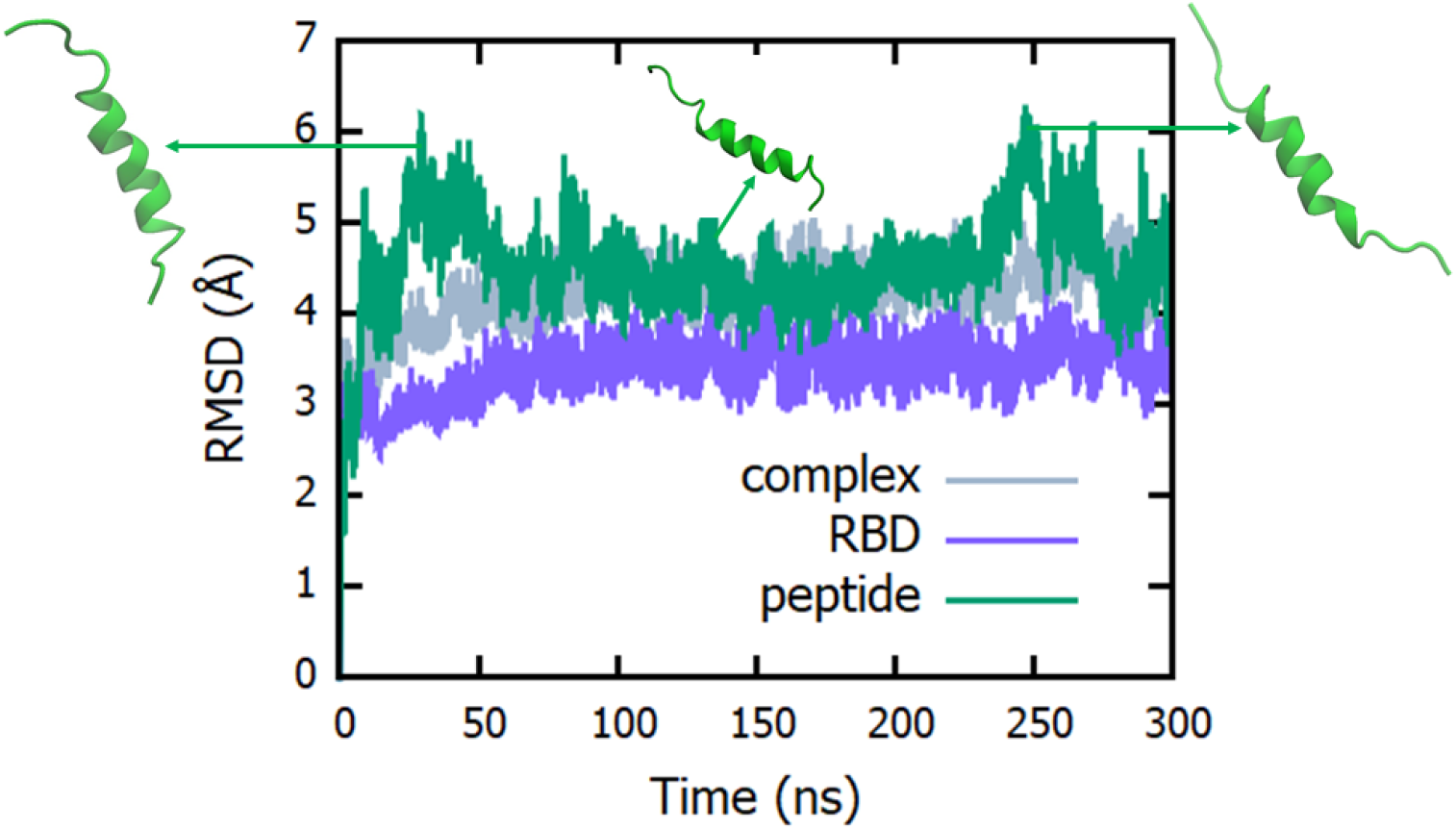
RMSD of the RBD-peptide complex as a function of time. Most of the contribution to the RMSD of the complex comes from the structural changes in the peptide. One notices a significant disruption of the peptide from its *α*-helical structure through the course of the simulation. Most of the disruption is in the terminal regions while the central region stays less disrupted.

**Figure 3:**
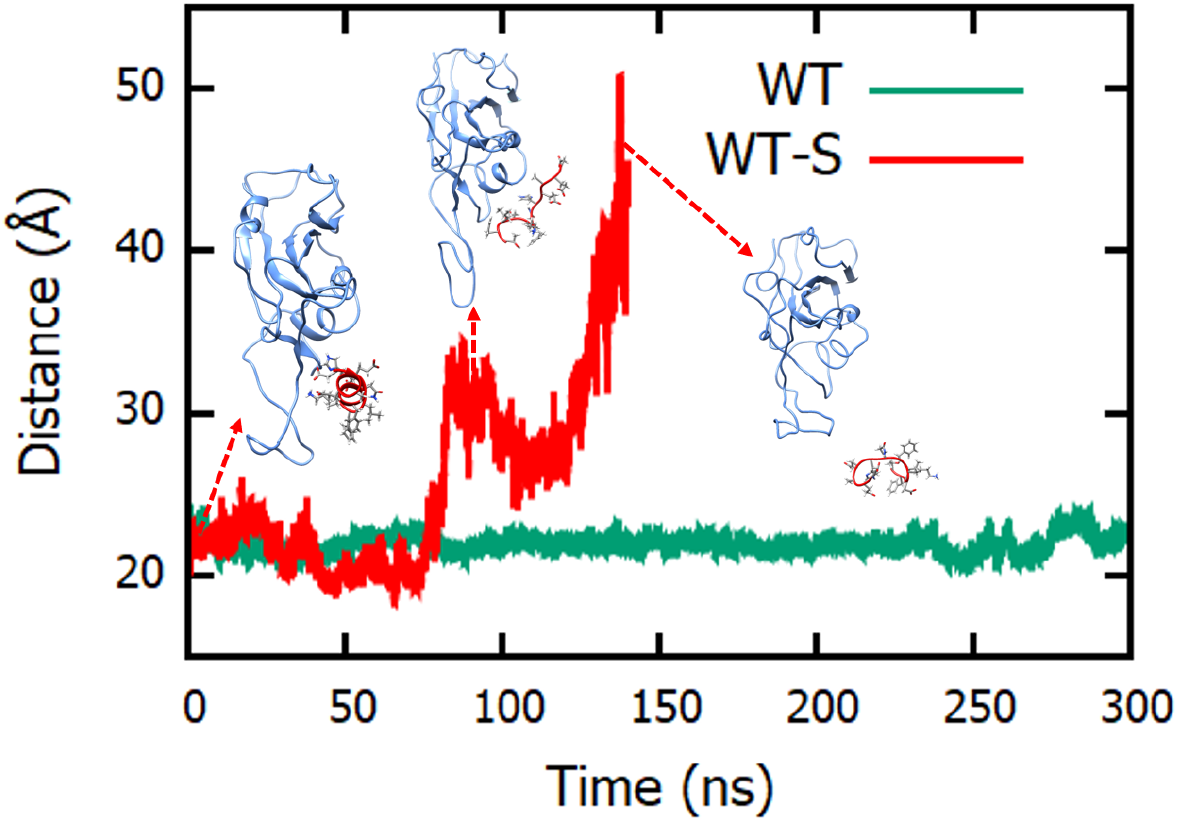
Distance between the centers of mass of the RBD and the peptide for the WT and WT-S systems as a function of time. One notices that while the full length peptide holds on firmly to the RBD, the truncated peptide not only flys apart but also loses its secondary structure.

Zhang *et al.*,^11^ from their RMSD calculations, found that the central 12-residue region of the peptide shows lesser fluctuations as compared to the terminal regions of the peptide in the peptide-RBD complex. From their residue-level RMSD calculations they infer that the region that shows lower RMSD (residue Thr7 to Asp18) may incorporate key contacts with the RBD. Our RMSD calculation shows that the majority of the contribution to the RMSD comes from the peptide. On observing the snapshots from the WT simulation we find significant disruption of the *α*-helical structure of the peptide at the termini, while the central region stays in tact (see fig.2), which would mean a lower RMSD for the central region. We think, that more than the presence of a majority of contacts in the central region of the peptide, this demonstrates significant dissolvability of the termini at the binding interface with the RBD, which suggests that the peptide termini are in an environment that provides strong interaction centers. Thus, in contrast to the inferences made by Zhang *et al.*,^11^ we feel that the terminal regions must also include a significant number of key interactions with the RBD.

For confirmation, we perform MMGBSA calculations on the peptide-RBD complex and find the residue-wise contribution to the binding energy for all the peptide residues (see fig.4). We observe that in addition to the central region of the peptide, the terminal regions (Ile1 to Lys6 and Leu19 to Ser23) also contain residues important for complex formation. Gln4 (−3.05 ± 1.5 kcal/mol), Ala5 (−1.4 ± 0.14 kcal/mol), Tyr21 (−4.0 ± 4 kcal/mol) and Gln22 (−2.0 ± 0.9 kcal/mol) have significant contribution to the binding energy. The contribution from the non-terminal region (Thr7 to Asp18) is dominated by residues Thr7 (−2.1 ± 0.4 kcal/mol), Phe8 (−2.6 ± 0.2 kcal/mol), Lys11 (−1.8 ± 0.3 kcal/mol), His14 (−2.0 ± 0.6 kcal/mol) and Asp18 (−1.5 ± 1.0 kcal/mol) (see fig.4).

**Figure 4:**
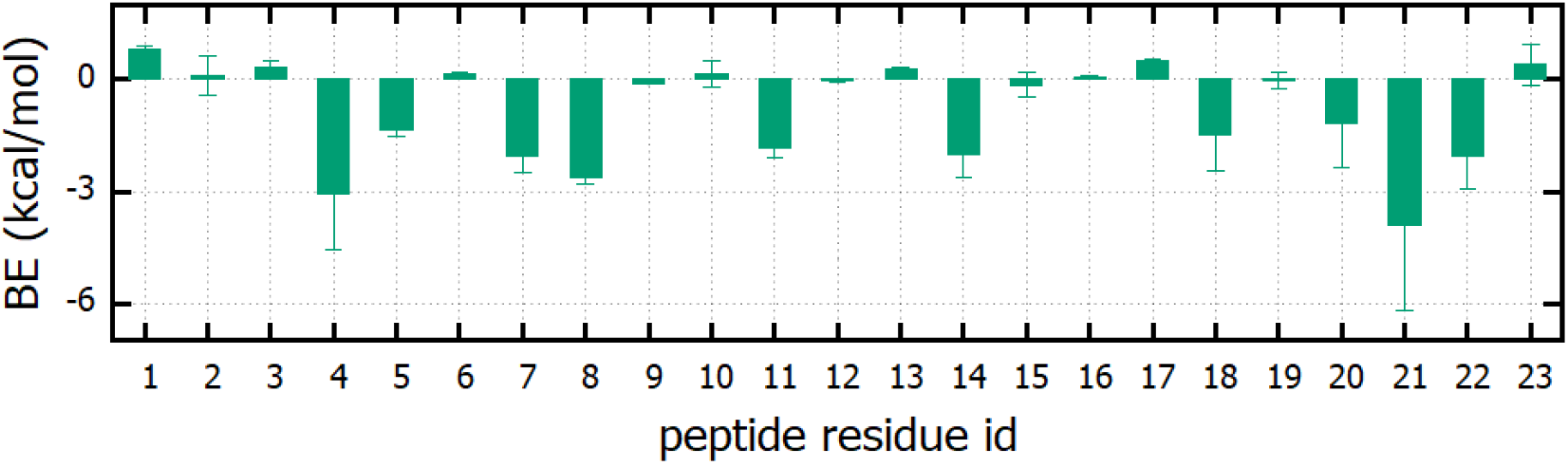
The residue-wise contribution of the peptide residues to the binding between the peptide and the RBD for the WT system. The average values have been calculated by first finding averages over five 10 ns windows from the last 100 ns of the MD trajectory and then taking the average over the five average values. The error bars are the standard deviations over the five averages.

From the residue-wise binding energy calculation we also find that the region between Leu9 and Phe20 contains many residues that do not contribute much to the overall binding energy. One can then modify these regions of low contribution in a way that the overall *α*-helical nature of the peptide is conserved, which will lead to a peptide variant that binds more efficiently to the RBD. The modified peptide would be able to engage in more contacts with the RBD, enhancing its viral blocking capability.

To test this, we performed simulations of various mutated peptides in complex with the viral RBD. The amino acid residues to be mutated was decided upon after an observation of the RBD-ACE2 binding interface in the 6M17^6^ structure and the peptide-RBD binding interface from the WT simulation. From figure 4, we observe that the lowest contribution to binding energy from the central region of the peptide comes from residues like Asn13, Glu15, Ala16, Glu17. We observe that, in the 6M17^6^ structure, Glu17 and Asp18 are in close proximity of two different Tyr residues of the RBD at the binding interface (see fig.S1 of Supplementary Information (SI)). As Glu17 does not form any contact with the RBD in the peptide-RBD complex, we decided to replace Glu17 to Leu giving rise to the E17L system. We wish to understand if the Leu residue would engage with the proximal Tyr residue forming strong hydrophobic contacts leading to a significant stabilizing contribution to the binding energy. Asp18 on the other hand has a significant contribution to the binding energy coming from a salt bridge that it forms with an Arg residue on the RBD. The interaction energy of the pair is ~−13 kcal/mol. In spite of forming such a strong salt bridge, the overall contribution of Asp18 to the binding energy is just ~−2 kcal/mol because of a strongly positive polar solvation energy of binding. We thus decided to mutate Asp18 to Leu (system D18L). This mutation would create the possibility of an interaction between the Leu residue and the Tyr that lies in close proximity to Asp18. On top of that we would be able to understand the position dependence of the effect of two equivalent mutations (a negatively charged residue to a hydrophobic residue) on the binding energy. Leu was selected as the mutation partner because it would preserve the *α*-helical structure of the peptide.^21^ Also, to understand the effect of multiple mutations present at the same time, we replaced Asn13 by Glu in E17L, giving rise to the system system N13E-E17L. This mutation is expected also to disrupt the *α*-helix because of a local accumulation of negative charge in the sequence and will give us an idea of the importance of rigidity of the peptide for an optimal binding to the RBD.

### E17L shows stronger binding to the RBD

From the calculation of the total energy of binding between the peptide and the RBD, we find that the E17L system shows a significant increase in the binding energy as compared to WT. Figure 5 shows a comparison between the peptide-RBD binding energies for the last 100 ns of the trajectory for the WT and E17L systems. While the average binding energy for the WT system is around −49 kcal/mol, the average value for the E17L system is about −63 kcal/mol.

**Figure 5:**
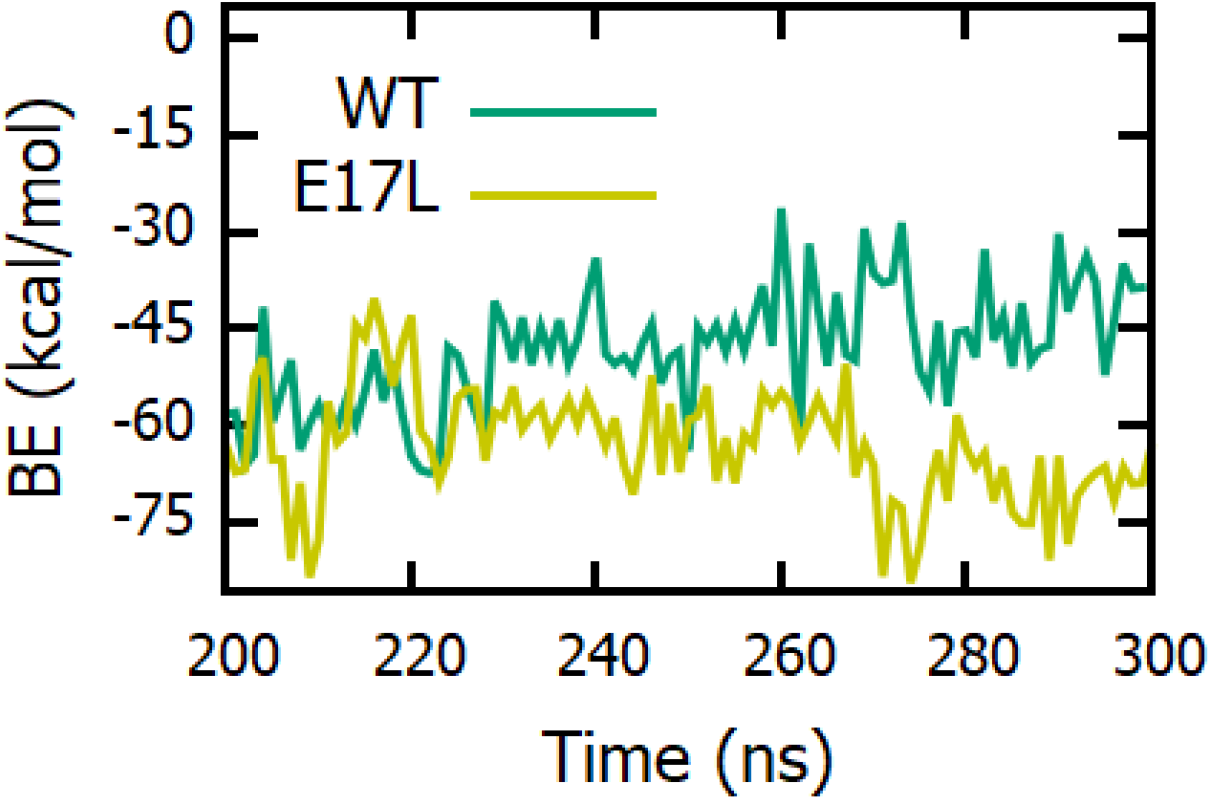
A comparison between the peptide-RBD binding energies for the WT and the E17L systems for the last 100 ns of the 300 ns long MD trajectory.

To obtain a deeper insight into this behaviour, we compare the residue-wise data for the two systems (fig 6). There are significant changes in the binding energy and contact distributions among different residues. In the E17L system, Leu17 (The mutation partner of Glu17 of the WT system) forms contacts with the RBD, whereas Glu17 was not forming any contacts in the WT system. While Glu17 had a slightly positive contribution to the peptide-RBD binding energy, Leu17 in the mutated system has a significant attractive contribution of ~−3.5 kcal/mol. In addition, many residues that were not forming any contacts with the RBD in the WT system, form significant number of contacts in case of E17L. Among them the most notable are Glu3, Asn13 and Phe20. Asp10 also shows a steep increase in the number of contacts. While there is an overall increase in the number of contacts, some other residues like Asp18, Phe8 and Glu15 show a reduction in the number of contacts with the RBD. To obtain a molecular picture and determine the origin of this behaviour, we visualize the binding interface (see fig.7) between the peptide and the RBD for the WT and E17L systems. As the contact plots suggest (fig.6), we observe that the interface is formed by different sets of peptide residues in the two systems. Guided by the strong hydrophopic contact between Leu17 and the Tyr residue which the incorporated mutation was targeted to engage, the orientation of the peptide, and thus the interface residues are completely different. With Leu17 forming contacts with the Tyr residue (see fig.7), the helical shape of the peptide and the resulting directionality of the amino acid side chains leads to a cooperative inclusion of residues like Asn13, Asp10, Phe20 and Glu3 to the binding interface. The helical structure ensures that only particular sets of peptide residues can form simultaneous contacts with the RBD. The total binding energy thus depends on which set of residues provide a more stable binding. Changing Glu17 to Leu shifts the balance in favor of a particular set of residues leading to larger stabilization of the complex.

**Figure 6:**
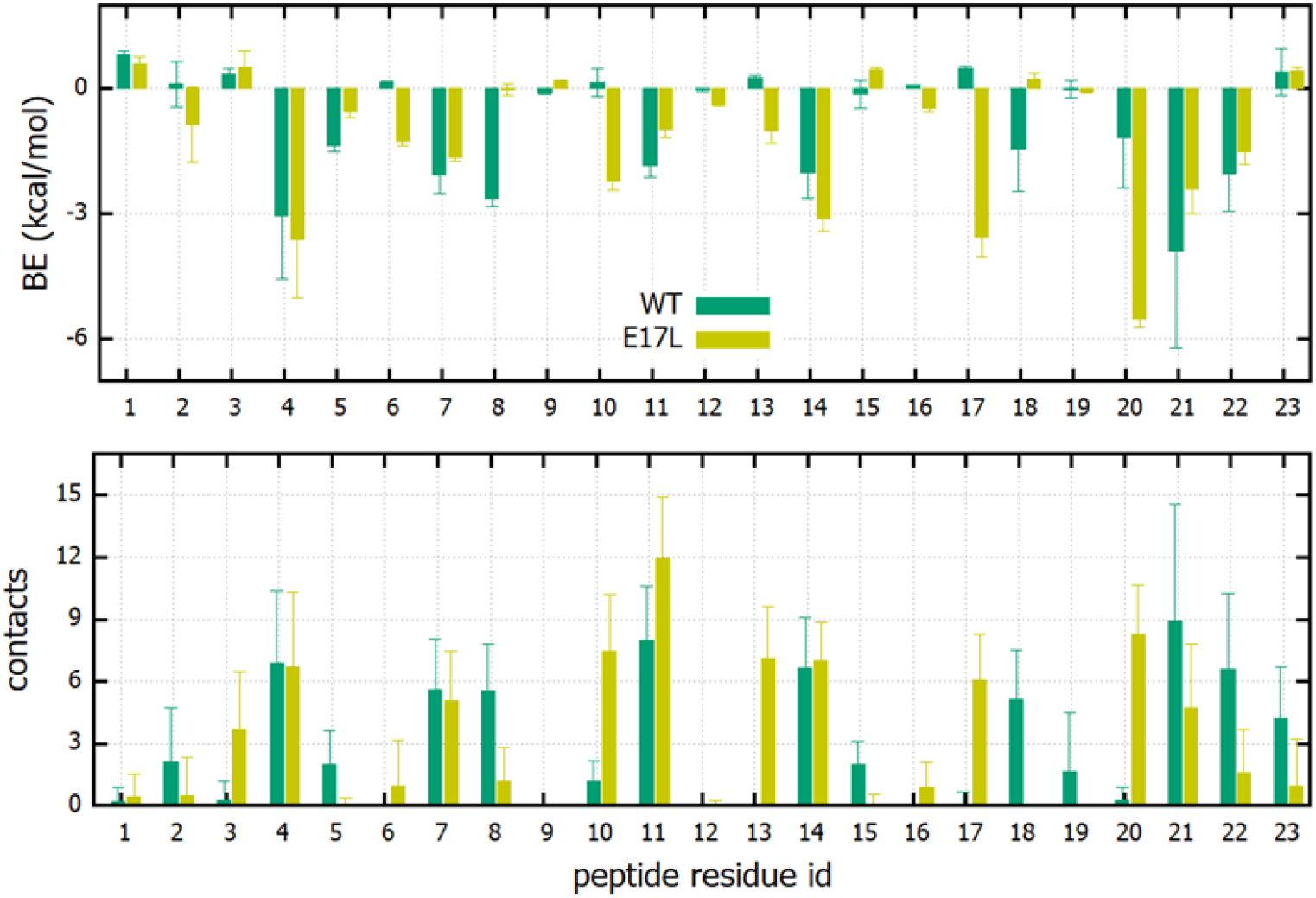
A residue-wise comparison between the peptide-RBD binding energy (top) and number of peptide-RBD contacts (bottom) for the WT and the E17L systems. Binding energy averages were first calculated over five 10 ns windows from the 200–300 ns section of the MD trajectory. The averages and error bars shown are determined over these five average values. The number of contacts are averaged over 1000 equally spaced frames from the last 100 ns of the MD trajectory.

**Figure 7:**
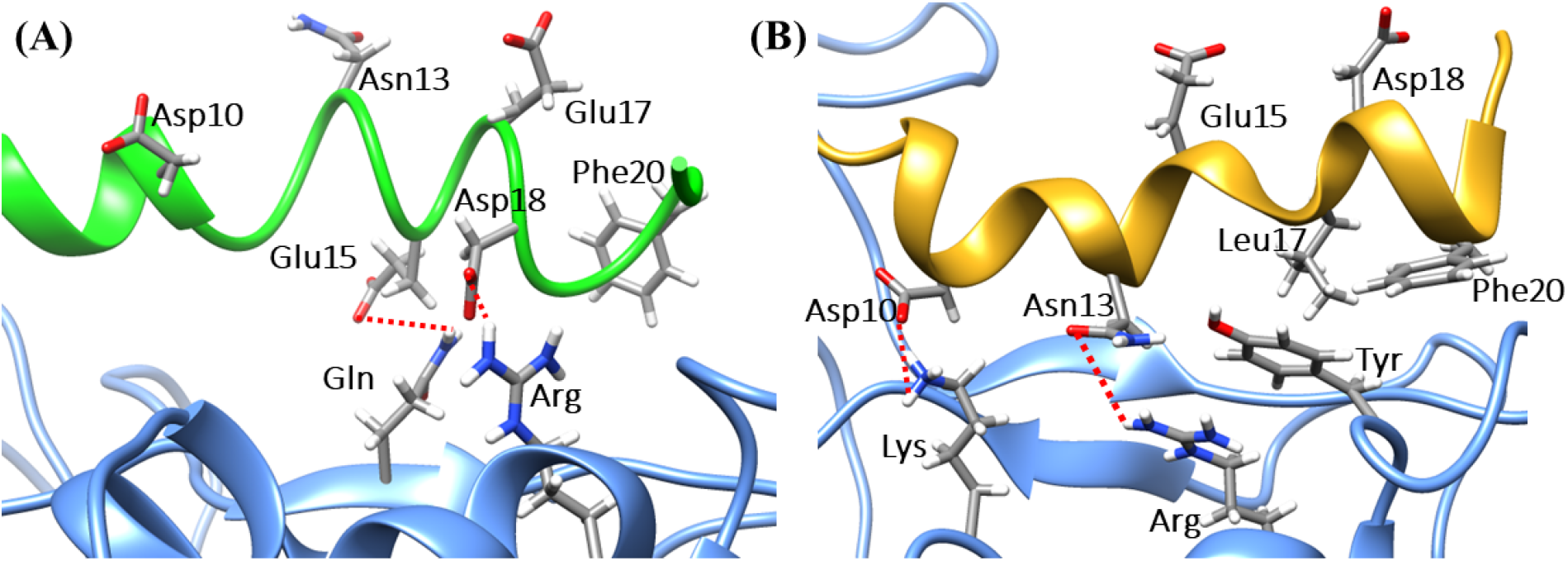
Snapshots depicting the residues that lie at the peptide-RBD binding interface for the (A) WT and (B) E17L systems. The RBD is shown in blue. The WT peptide is shown in green while the E17L variant is shown in golden color.

**Figure 8:**
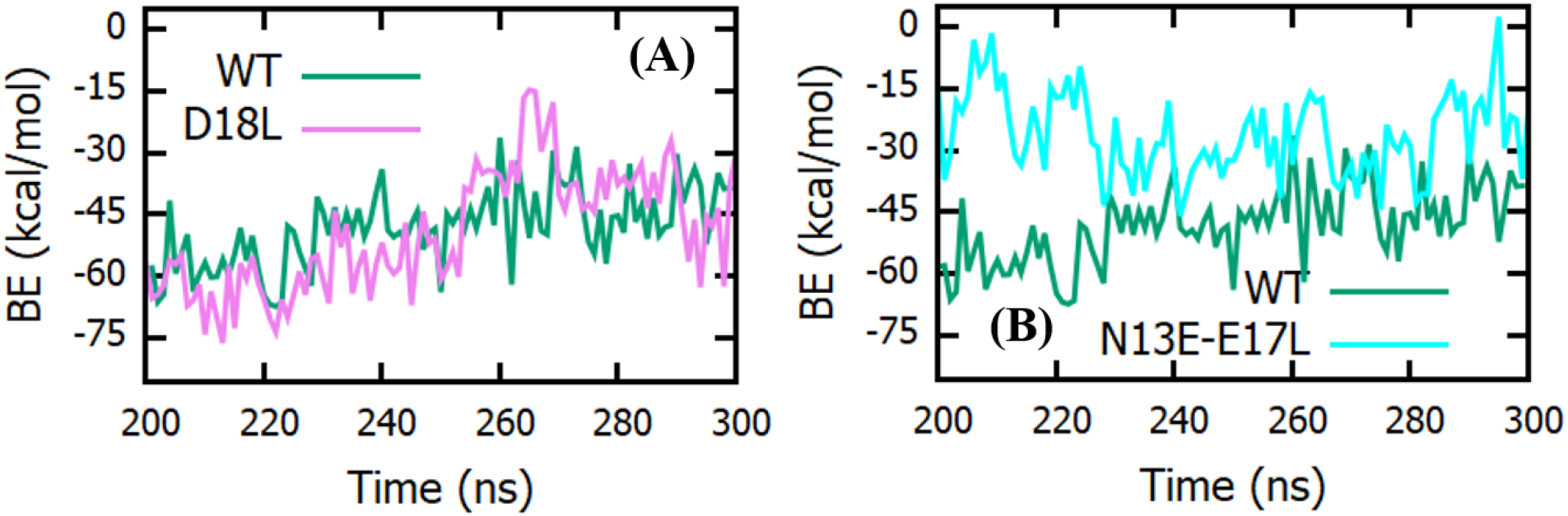
A comparison of the peptide-RBD binding energies for the (A) D18L and (B) N13E-E17L systems with the WT system for the last 100ns of the MD trajectory.

### The α-helical structure is crucial for optimal binding

While we observe that the mutation E17L leads to a significant stabilization of the peptide-RBD complex, we observe very different results for the D18 system. The average binding energy for the D18L system was calculated to be ~−50 kcal/mol. This was almost similar to the value of ~−49 kcal/mol for the WT system. The snapshots in fig.7 provide some hint towards this behaviour. As Asp18 was already forming contacts with the RBD in the WT simulation, replacing it with a Leu would lead to a binding interface that must be very simular to what one observes for the WT system. This also shows that the stabilizing effect of the E17L mutation was not just a result of an increase in hydrophobicity of the peptide but was a combination of hydrophobic effect and the choice of the mutation point. Although, Asp18 lies just next to Glu17 in the peptide sequence, but as discussed above, the directionality in interactions introduced by the helical nature of the peptide leads to a significant difference between the effects of the two mutations. Thus the helical structure dictates the behaviour of the peptide-RBD energetics.

One might then wonder what happens if mutations that can disrupt the helical structure of the peptide are incorporated. The case of N13E-E17L is hence of interest. In the E17L simulation we notice that Asn13 was forming contacts with an Arg residue of the RBD (see fig.7), which we expected to engage in a strong salt-bridge with the Glu that replaces Asn13. On the contrary the mutation led to a disruption of the peptide helix (see fig.S2 of SI) because of intra-molecular electrostatic strain as a result of the presence of Asp10 in close proximity (see fig.7). As a result of the disruption of the helix and the peptide losing structure and rigidity, we observe a strong reduction in binding energy. In the absence of other stabilizing interactions, even Leu17 did not form the hydrophobic contacts that it took part in during the E17L simulation. The average binding energy for the N13E-E17L system is around 26 kcal/mol. We calculated the distance between the centers of mass of the peptide and the RBD for all the modified peptide systems (see fig.S3 of SI). We observed that, although the peptide in case of N13E-E17L stays attached to the RBD throughout the simulation, it shows enhanced fluctuations not present in other systems. An unstructured peptide would introduce strong entropic factors that won’t favor a bound state, something that we observed in case of the WT-S simulation. Thus, multiple mutations (or a single mutation for that matter) should be chosen carefully such that the helical nature of the peptide stays intact. This shows that the helix not only aids in specificity via a proper recognition of the binding ridge but also is crucial in ensuring an optimal binding.

The average binding energies of all the systems are summarised in table 2.

**Table 2:** Peptide-RBD binding energies for different systems.

| Complex | Binding energy (kcal/mol)<br>(whole trajectory) | Binding energy (kcal/mol)<br>(last 100 ns) |
| --- | --- | --- |
| Wild type | $-48.8 \pm 9.9$ | $-48.4 \pm 9$ |
| E17L | $-57.0 \pm 10$ | $-63.0 \pm 8.8$ |
| D18L | $-50.0 \pm 12$ | $-49.5 \pm 14$ |
| N13E-E17L | $-26.3 \pm 10$ | $-26.3 \pm 10$ |

## Conclusion

In this work, we have performed MD simulation of a complex between the SARS-CoV2 RBD and a peptide fragment that is a part of the viral binding region of ACE2. The peptide has been recently proposed as a potential drug candidate against Covid-19. From our simulations of the wild type peptide in complex with the RBD, we found, contrary to previous results, that the residues that form key contacts with the RBD are well distributed throughout the length of the peptide abd are not confned to the central region of the peptide. We determined the residue-level contribution to binding energy of the peptide-RBD complex. This helped us to single out residues that have minimal contribution to the binding energy. We modelled mutations in the peptide at points of low contribution to the binding energy and demonstrated that the peptide variant E17L bound stronger to the RBD as compared to the wild type peptide. Recognizing such modifications that would preserve the *α*-helical nature of the peptide while increasing the affinity towards the target is crucial in rendering the peptide more effective in viral blocking. We also find that the residues bind cooperatively with the RBD. The cooperativity can be positive or negative and has its roots in the helical structure of the peptide that introduces directional effects. We also investigated the importance of the structure and rigidity of the peptide and found that mutations that disrupt the rigid helical structure of the peptide lead to a very weak binding to the RBD. Our results have completely been derived using computational methods but can be easily tested by experiments.

The strategy of using human-derived peptides as drugs is attractive, but one needs to recognise that the peptides that are supposed to hinder a particular biological pathway, of which they themselves are the central player as a part of a larger globular protein, will be prone to competitive replacement. Modifications that have a stabilizing effect on the peptide-target complex are thus crucial for the proper utilization of drug-like properties of those peptides which are supposed to mimic themselves. One can thus utilise their modifiability and tune them in a target-specific manner to positively affect their performance.

## Supplementary Information

**Figure S1:**
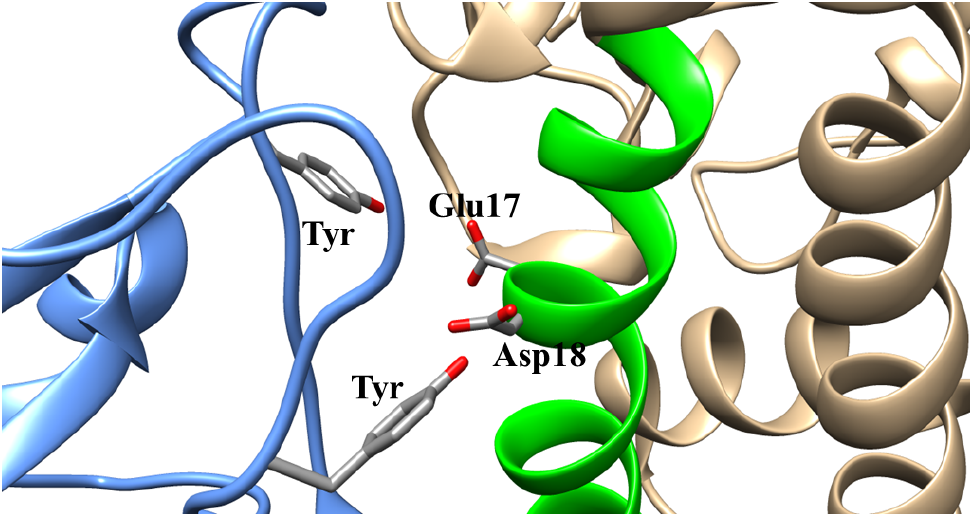
The ACE2-RBD interface. The 23-residue peptide is shown in green. The RBD is shown in blue. One can see Glu17 and Asp18 of the peptide pointing towards two Tyr residues on the RBD.

**Figure S2:**
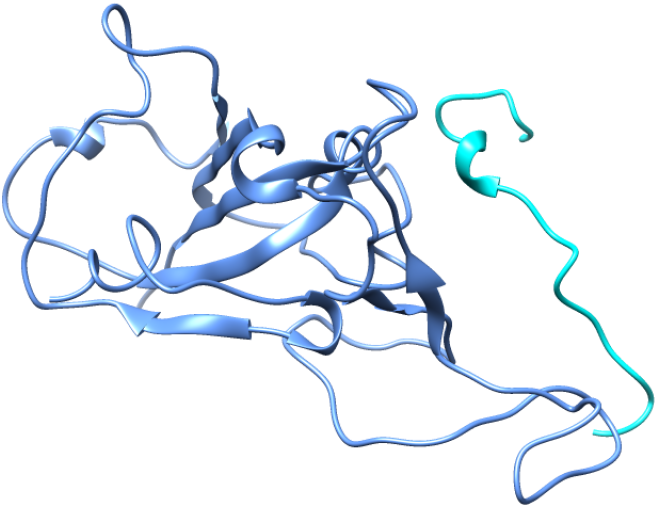
The peptide-RBD complex for the N13E-E17L system. The peptide is shown in cyan and the RBD is shown in blue. One can clearly see that the peptide has lost its helical structure and turned unstructured.

**Figure S3:**
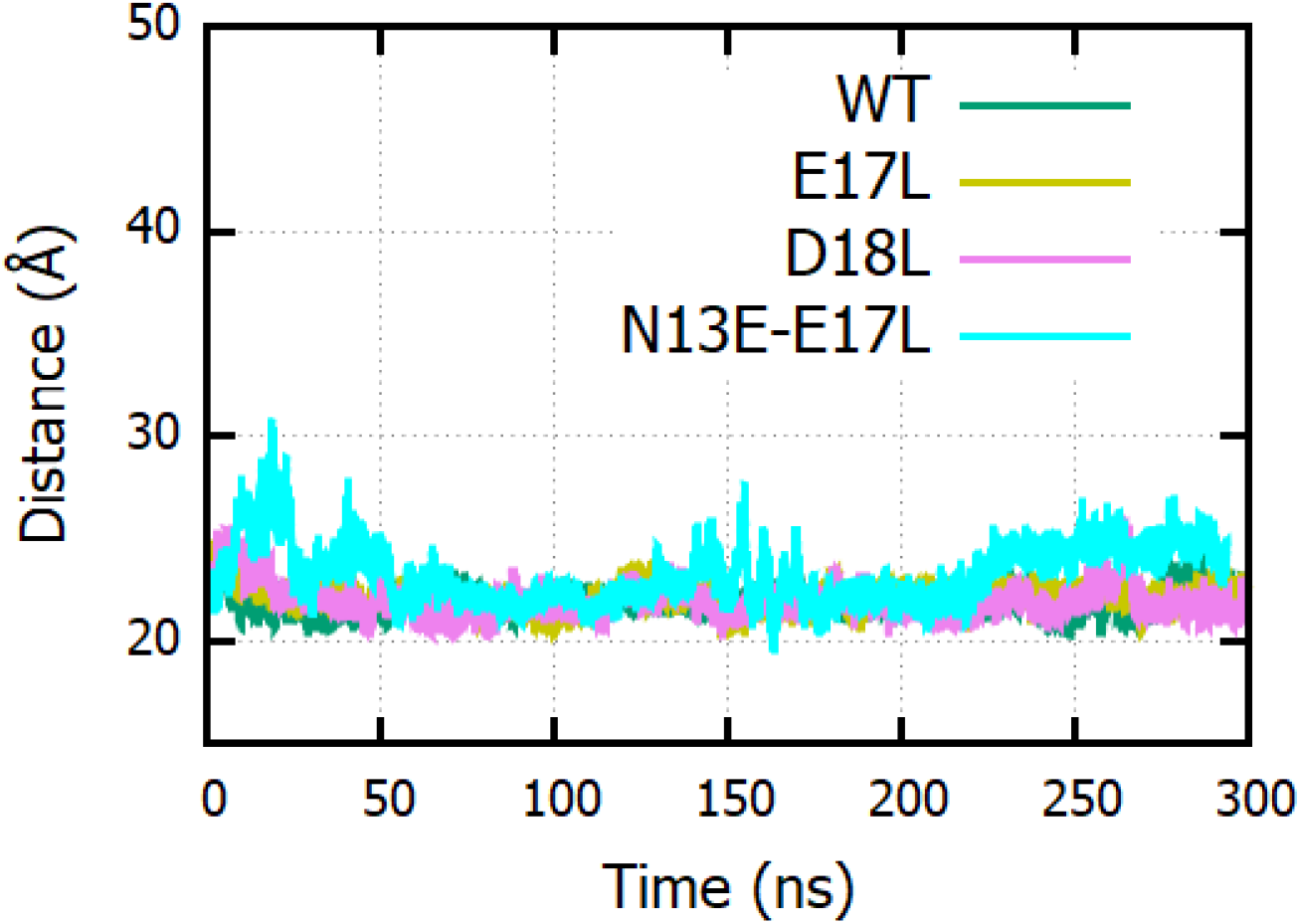
The distance between the centres of mass of the peptide and the RBD for the different full-length peptide systems simulated in this work. One observes enhanced fluctuations for the N13E-E17L system.

